# Performance Assessment of an Unsupervised Variable Selection Approach for Biomarker Discovery and Glioblastoma Subtyping

**DOI:** 10.1101/2025.03.11.642670

**Authors:** Roberta Coletti, J. Orestes Cerdeira, Marcos Raydan, Marta B. Lopes

## Abstract

High-dimensional omics data often contain more variables than observations, which negatively impacts the performance of classical data analysis methods. Dimensionality reduction is typically addressed through variable selection strategies that incorporate a penalty term into the model. While effective for selecting task-specific variables, this approach may not be optimal when the goal is to preserve the dataset structure and the overall biological information for multiple downstream analyses. In such cases, a priori unsupervised variable selection is preferable. In this study, we evaluate several unsupervised variable selection approaches to derive a representative subset of the original dataset. Building on the performance assessment results, we introduce TRIM-IT, a novel tool for unsupervised variable selection, clustering, survival analysis, and differential gene expression analysis. Applied to glioblastoma (GBM) data, TRIM-IT identified three clusters that correlate with tumor histology, exhibit distinct survival curves, and display unique molecular profiles with genes potentially serving as biomarkers. The tool is available for reproduction and adaptation to other studies.

## 1. Introduction

Variable selection has become a critical topic in modern data analysis due to rapid technological advances that enable the collection of large-scale, high-dimensional data. These datasets often exhibit more variables than observations, rendering many traditional statistical methods inadequate or computationally unfeasible. In this context, variable selection methods aim at reducing the dimensionality of data sets while minimizing the loss of information; see, e.g., [1, 2, 3, 4, 5, 6, 7, 8].

In many cases, variable selection is integrated into a classical statistical model by adding a regularization term, such as the *ℓ*_1_ norm. While this approach is valuable when the goal is to examine data for a specific purpose (e.g., classification or survival analysis), it is less suitable when more comprehensive analyses would be performed on the same set of variables. In these cases, the objective should be to identify, by an unsupervised approach, a subset of variables that closely approximates the complete dataset, to be analyzed from different perspectives, with classical methods and without loss of generality.

Unsupervised variable selection is a challenging task. Most of the criteria found in the literature are focused on ranking variables based on their relevance, which can be measured in different ways. For instance, it is known that variables with no variability along the different observations do not carry much information [9]. Therefore, the Variance Criterion (VC) ranks the variables based on their variance. Similarly, the relevance of a variable is also associated to its dispersion, which can be computed by the mean absolute difference (MAD), measuring the variable’s deviation from its mean. However, this approach does not take into account redundancy among variables, which might negatively affect the results; if two similar variables are relevant, including both in the selection does not increase much the overall information. Therefore, another strategy is based on filtering variables based on both relevance and redundancy (RR) [10]. The RR algorithm starts by considering the first variable in terms of relevance, then the following one is taken only if it is not similar to the variable/s already selected, based on a predefined threshold. A different approach to rank variables is by considering the Laplacian Score (LS), a graph-based method that focuses on geometrical properties. LS quantifies the importance of a variable based on its ability to preserving the local data structure [11].

Ideally, the subset of variables selected by an unsupervised approach should be representative of the original data. Many criteria have been proposed to assess how well a subset of features approximates the whole dataset [1, 12, 13]. The basic idea is to consider the dataset *X* as a *n* × *p* matrix, where *n* is the number of observations and *p* the number of variables, and to compute the similarity between *X* and *X*^*r*^, the latter representing a reduced dataset with *r* variables, with *r < p*. Depending on the data property to preserve, different coefficients might be defined. The RM coefficient gives an indication of the similarity between the spectral decomposition of the full and the reduced data matrices [1]. It measures the cosine of the angle between the full data matrix *X* and the one resulting from the projection of *X* on the *r*-dimensional subspace generated from the subset of *r* variables. The RV coefficient [14] is based on preserving the geometrical representation of two configurations of *n* points (*X* and *X*^*r*^), quantifying their similarity in a linear space. Finally, the GCD coefficient is based on the principal component analysis, and it measures the distance between *X*^*r*^ and the subspace given by the first *r* components of the original space *X* [1].

Finding the best subset based on a given criterion can be challenging, and optimization approaches may be employed to achieve this goal. Among them, simulating annealing [15] is a general probabilistic optimization procedure that at each step randomly selects some neighbor solution *S*’ of the current feasible solution *S*, and probabilistically decides whether to keep *S* as current or to replace *S* by *S*’. A simulated annealing algorithm (anneal) designed to optimize each of the three criteria: RM, RV and GCD, is proposed in [12].

The different variable selection approaches usually need to set certain parameters that affect the final results. Moreover, most of these methods require that the cardinality of the desired subset is given in advance. Choosing the right cardinality is not always straightforward, and may result in suboptimal solutions [16]. To address these limitations, we considered the Max-Out Min-In Problem (MOMIP), a graph-based optimization method designed for variable selection. The great advantage of this method is that it does not require knowing in advance the number of variables to provide an optimal solution, which exclusively depends on the data structure.

In this work, we aim to perform unsupervised variable selection in glioma gene expression data. Gliomas are a family of brain tumors. The high heterogeneity characterizing this cancer is one of the main reasons for treatment failures and poor overall patients’ survival. The recent advances in the molecular understanding of gliomas allowed for the definition of three main cancer types by the World Health Organization, namely astrocytoma, and oligodendroglioma, also designated Low-Grade Gliomas (LGG) and glioblastoma (GBM), which is the most aggressive type. However, studies revealed that GBM patients show diverse survivals and treatment responses, suggesting the existence of further patients’ stratification [17, 18, 19]. Despite recent efforts in this direction, the characteristics of such subtypes are still not fully elucidated [20, 21, 22]. In this context, unsupervised approaches for variable selection and sample grouping are particularly suitable to increase our understanding of GBM.

Clustering methods allow for unsupervised patients’ stratification solely based on their molecular profiles. In recent years, many clustering methodologies have been proposed, but they might fail when dealing with highdimensional data, which considerably increases the computational cost and compromises the outcomes [23, 24]. From this perspective, a prior variable selection highly increases the reliability of clustering results and improves the interpretability.

This study is divided into two stages. The first one was devoted to determining the unsupervised variable selection method that is more suitable for glioma data. In this phase, the complete set of data was used, as we employed prior clinical knowledge of glioma classification to evaluate the goodness of the selected subset of variables. Ideally, as we aimed to reduce as much as possible the dependency on the parameter setting, MOMIP would be the preferred method, but its outcomes were compared to other well-known unsupervised approaches. The method performances were evaluated in terms of the ability of the subset of variables to represent the full dataset by RM, RV and GCD coefficients. Moreover, we would ensure that the process of variable reduction preserves the intrinsic differences among glioma types. To evaluate this, unsupervised clustering was applied to the full and reduced dataset, and the obtained clusters were compared with the known glioma groups.

The second stage was focused on developing TRIM-IT, a novel tool that begins by applying the selected unsupervised variable selection method to GBM data. Beyond identifying significant features, this dimensionality reduction step addresses the imbalance between the number of samples and variables, which is critical for downstream analyses. Specifically, TRIM-IT aims to uncover robust GBM subtypes that may be linked to distinct patient characteristics or variations in life expectancy.

The paper is structured as follows. Section 2 presents the materials and methods, detailing the data and methodology used in our study. Section 3 reports the results, with a discussion of the findings. Finally, Section 4 concludes the paper with a summary of key points, highlighting the strengths and limitations of our work.

## 2. Materials and methods

### 2.1. Data

The glioma RNA-sequencing (RNA-Seq) dataset used for this work was collected from The Cancer Genome Atlas (TCGA) program, under the LGG and GBM projects [25, 26, 27]. In particular, the data were downloaded from the GDAC Broad Firehose (https://gdac.broadinstitute.org) portal, which aggregates TCGA information to be easily retrieved in a matrix form. In this study, the number of variables of the full RNA-Seq dataset was previously reduced from 20K to 145 variables, focusing on the genes coding for proteins selected by The Cancer Proteome Atlas database to be involved in important biochemical pathways in cancer [28]. This filtering step improved dataset manageability and the computational efficiency of subsequent algorithms, while ensuring that variables crucial for major cancer-related processes were maintained in the analysis.

The TCGA RNA-Seq data are already normalized by upper quartile normalization, to minimize variability of data collected by different institutions. To apply unsupervised variable selection and clustering, we considered RNA-Seq data treated by TCGA with Transcript Per Million (TPM) normalization, which takes into account gene lengths and sequencing depth. Additionally, the data were normalized by the log-2 transformation, a standard approach to RNA-Seq data as it preserves the biologically-detectable changes while stabilizing variance and enhancing comparability [29]. Conversely, to perform the differential gene expression analysis raw counts data were used.

To evaluate the clustering results, glioma samples were labeled with the corresponding glioma type updated to the 2021-WHO classification guidelines [30, 31], resulting in 206 astrocytomas, 143 oligodendrogliomas, and 116 GBM samples. See the data availability section for further details.

### 2.2. Analysis workflow

This work is structured into two phases: (1) an exploratory stage, in which MOMIP was compared with other well-known methods for unsupervised variable selection to evaluate their performances on glioma data, and (2) the application of the method identified as the most suitable for our goal to a glioma case of study by defining TRIM-IT.

In the first phase, the whole glioma dataset comprising of 465 patients and 145 variables was considered. The MOMIP’s performance was compared with seven other unsupervised variable selection approaches: VC, LS, MAD, RR filter, and the anneal algorithm applied to RV, RM and GCD criteria. As MOMIP is not regulated by arbitrary parameters, we first applied MOMIP method to glioma data, and then the other algorithm/criteria were appropriately set to select the same number of variables, allowing performance comparison. MOMIP’s lack of parameter dependency can be a limitation if the final number of selected variables is larger than desired. In such cases, MOMIP can be applied recursively until the target subset size is achieved. In this study, MOMIP was employed in two recursive steps: first reducing the variable count from 145 to 49, and then further down to 18. Therefore, the other methods were set accordingly: anneal was run with a predefined number of 18 features to select, while VC, LS, and MAD were used to rank the variables and then select the top 18. The RR filter was applied with a similarity threshold corresponding to the maximum value that allowed for the selection of 18 variables, ensuring the removal of redundant features.

For each method, the subset of selected variables was evaluated in terms of its representability of the original dataset – by RV, RM and GCD coefficients– and based on the ability to separating the known glioma types by K-means clustering. Clustering results were assessed by supervised (Adjusted Rand Index, Adjusted Mutual Information, and Normalized Mutual Information) and unsupervised measures (silhouette and Calinski-Harabasz scores).

To validate the robustness of the results, we also compared them with those obtained from 1000 random subsets of 18 variables from the original dataset. This comparison helped rule out the possibility that the observed results were due to chance, confirming that the selected variable subsets provided meaningful insights rather than random patterns.

In the second part of this study, we propose TRIM-IT, a tool for unsupervised variable selection that ensures retention of overall data information for patient stratification. TRIM-IT is designed to first select informative variables trough MOMIP, and then analyze the reduced dataset from multiple perspectives, enabling a comprehensive study (scheme in Figure 1). We applied TRIM-IT to GBM data with the aim of identifying biologically meaningful GBM subtypes. The GBM dataset (116 observations for 145 variables) was reduced by performing variable selection through MOMIP (step 1, Figure 1), leading to a subset of 19 variables. Then, K-means clustering was applied to stratify patients (step 2, Figure 1). The identified clusters were analyzed to determine if the patients in each group shared specific characteristics. Clinical data were used to identify factors such as sex, histology, and survival that could correlate with the clusters (step 3, Figure 1). Additionally, Differential Gene Expression (DGE) analysis was conducted to further characterize the groups by gene expression (step 4, Figure 1). For technical details, see Supplementary Materials sections S2 and S3.

**Figure 1:**
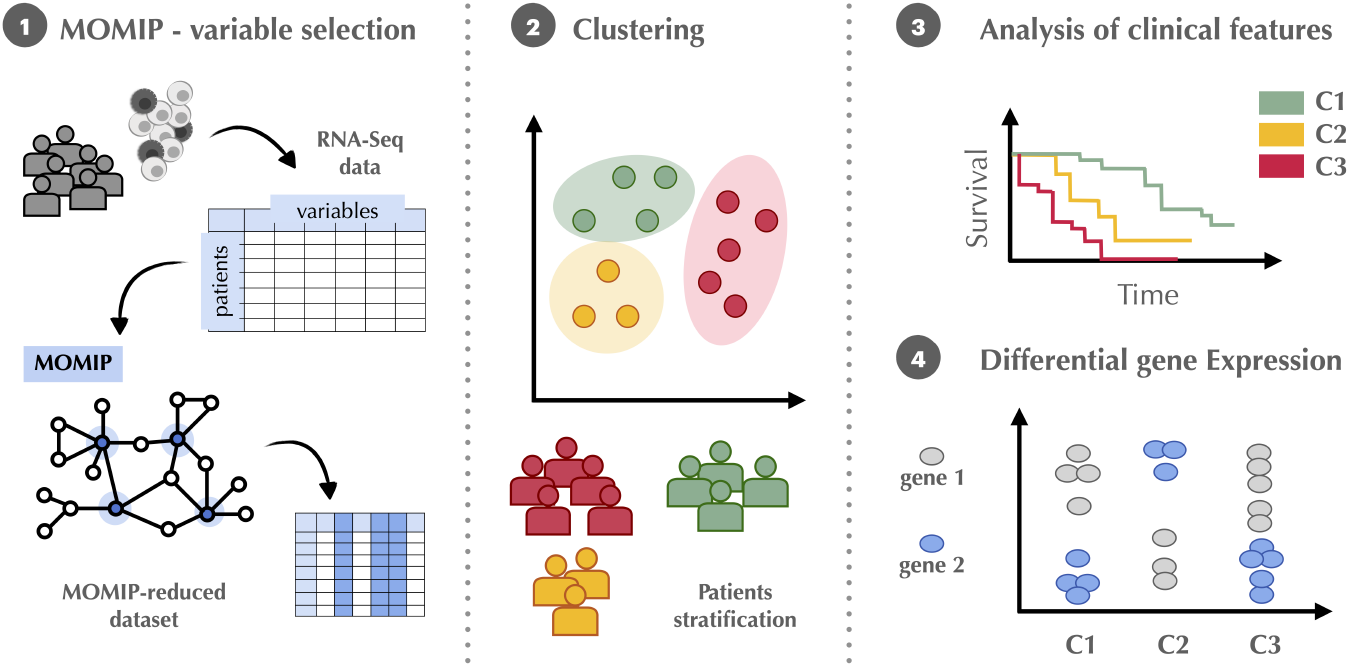
TRIM-IT scheme. Step 1: MOMIP - variable selection. Bulk RNA-sequencing data were retrieved from the TCGA portal in matrix form, with observations (GBM patients) as rows and variables (genes) as columns. MOMIP was used for unsupervised variable selection, reducing the original dataset of 145 variables to a refined subset of 19 key variables. Step 2: Clustering. Using the reduced dataset of 116 patients and 19 variables, K-means clustering was performed to stratify patients into three groups. Step 3: Analysis of clinical features. Clinical data were analyzed to identify features characterizing each cluster. Kaplan-Meier survival curves were also obtained to compare survival outcomes across the three patient groups. Step 4: Differential Gene Expression. The reduced RNA-sequencing dataset was analyzed to identify genes differentially expressed across the three clusters.

All the analyses shown in this study were performed in software R (version 4.3.2). To allow the reproducibility of results, the implemented R scripts and used data are available at: //github.com/RobertaColetti/TRIM-IT.

### 2.3. The max-out min-in problem

The max-out min-in problem (MOMIP) is a combinatorial optimization problem that adequately applies to variable selection [32].

Given a set of *p* entities and weights *w*_*ij*_ ≥ 0 between every two entities *i, j* ∈ *N*, with *w*_*ij*_ = *w*_*ji*_ and *w*_*ii*_ = 0, MOMIP looks for a subset *S* of entities that maximizes the sum of the weights of pairs of entities that have exactly one entity in *S*, while minimizes the sum of the weights between entities of *S*. Thus, MOMIP seeks at

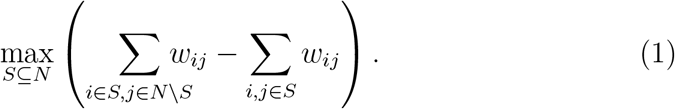

Suppose now that *X* is an *n* × *p* matrix of measurements of *p* variables (columns) in *n* objects (rows). If we let *N* to be the set of the *p* columns of *X* and *w*_*ij*_ the square of the Pearson correlation between the variables *i* and *j, i* ≠ *j* = 1,…, *p*, and *w*_*ii*_ = 0, MOMIP will select a set *S* of columns that are low correlated to each other and highly correlated to the columns that are not selected, i.e., a set of maximal *independent* variables. This procedure has the advantage of not requiring the number of variables to be known in advance.

### 2.4. Clustering

K-means clustering consists in dividing samples into clusters such that the distance of its sample to the cluster center (centroid) is minimized. Specifically, K-means clustering finds the best partition of the sample set *{x}* into *K* (nonempty) clusters *C*_1_, …, *C*_*K*_ by solving the following minimization problem:

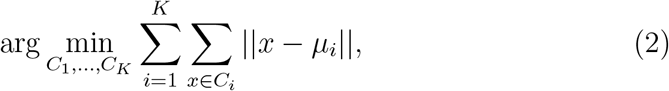

where 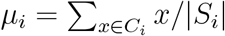 is the centroid of the cluster *i, C*_*i*_ ∩ *C*_*j*_ = Ø,*i* ≠ *j* = 1,…, *K*, and 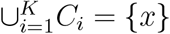.

Clustering performances are evaluated by using some measures, which can be distinguished in supervised and unsupervised. Supervised measures are used to assess the ability of the clustering method to recognize known groups, so they are employable in case patients’ labels are available. The most well-known supervised measure is the Rand Index (RI), which quantifies the similarity between clusters (*C*) and patients’ labels (*L*) by computing the ratio of correct assignments (true positive and true negative) to the total number of samples [33]. However, RI tends to be artificially high, as it does not account for the likelihood of some associations occurring by chance. Therefore, it is preferred considering the Adjusted Rand Index (ARI), which modifies the formula by introducing the expected RI value (𝔼 (*RI*(*C, L*))):

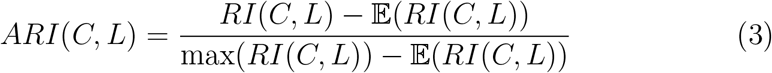

Differently from RI, which is between 0 and 1, ARI can reach also negative values. Random cluster assignments correspond to 0, while higher values mean a positive agreement between the two patients’ grouping. Other supervised measures are based on mutual information (MI), which quantifies the shared information between two variables. In the context of clustering, MI measures the reduction in entropy (*H*) of the clustering result (*C*) given knowledge of the true labels (*L*), i.e., *MI*(*C, L*) = *H*(*C*) — *H*(*C*|*L*). A high value of MI means that the clustering result aligns with the ground truth labels. To facilitate comparison among clustering results, Normalized Mutual Information (NMI) is often used; this scales MI values in [0, 1], where 0 means no agreement and 1 indicates perfect correlation:

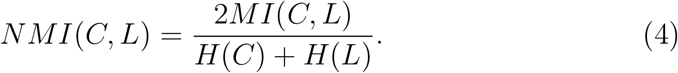

As for RI, MI does not consider the potential random assignments; the Adjusted Mutual Information (AMI) is defined by introducing the expected MI value:

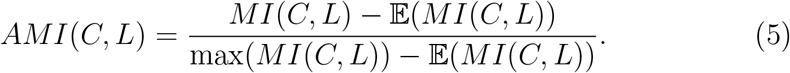

When the true labels are not available, unsupervised measures can be used to evaluate the quality of the cluster structure. The most known unsupervised score is the Silhouette (*s*), which measures the distance of each observation to the cluster to which it belongs, with respect to the nearest cluster. Let *a*(*i*) be the average distance of the sample *i* to all the samples of the same cluster, while *b*(*i*) the average distance of *i* to the samples of the nearest cluster. The *s* score of the sample *i* is:

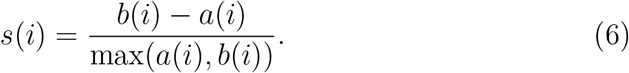

The *s* score ranges between −1 and 1, where higher values mean better defined and well-separated clusters, while a negative value suggest that the sample may be incorrectly assigned to its cluster. By assigning a score to each data point, silhouette measure quantifies how well a single sample fits with the assigned cluster. To assess the overall cluster quality, the average silhouette score is commonly used, though it may not fully capture the variability of individual point scores.

Another unsupervised measure is the Calinski-Harabasz (CH) score, which evaluates clusters’ structures based on the ratio of the sum of between-cluster dispersion to within-cluster dispersion [34]. Let *B* be the between-group dispersion matrix, and *W* the within-group dispersion matrix, the trace of *B, tr*(*B*), is the sum of squared distances between cluster centers and the global center, while the trace of *W* is the sum of squared distances within each cluster. Therefore, the CH is defined as:

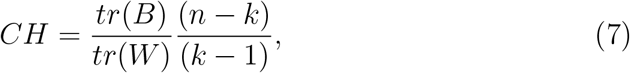

where *K* is the number of clusters and *n* the number of samples.

Higher CH scores indicate better-defined clusters, characterized by greater separation between clusters and tighter cohesion within them. Since the CH score does not have an upper bound, it must be compared across different results to be properly interpreted. Nevertheless, by providing an overall measure of cluster quality, the CH score is particularly useful for comparing the performance of different clustering algorithms.

## 3. Results and Discussion

This section is structured into two subsections: the first related to the exploration phase of our study, in which MOMIP performances are compared with other unsupervised methods for variable selection, while the second presents the results of TRIM-IT, where MOMIP and K-means clustering were applied to the GBM dataset.

### 3.1. Comparison of unsupervised variable selection methods

We compared the performance of eight unsupervised variable selection approaches based on their ability to select a subset of variables that effectively represents the entire starting dataset while preserving the biological information necessary to accurately cluster samples into known glioma types. To evaluate the representational power of these methods, we considered the three criteria introduced above: RM, RV, and GCD. Figure 2 reports the values of these coefficients for each subset identified by a given method (indicated by colored vertical lines). These results were also compared against a random baseline, which consists of the coefficients calculated for 1,000 random subsets of 18 variables, represented by histograms.

**Figure 2:**
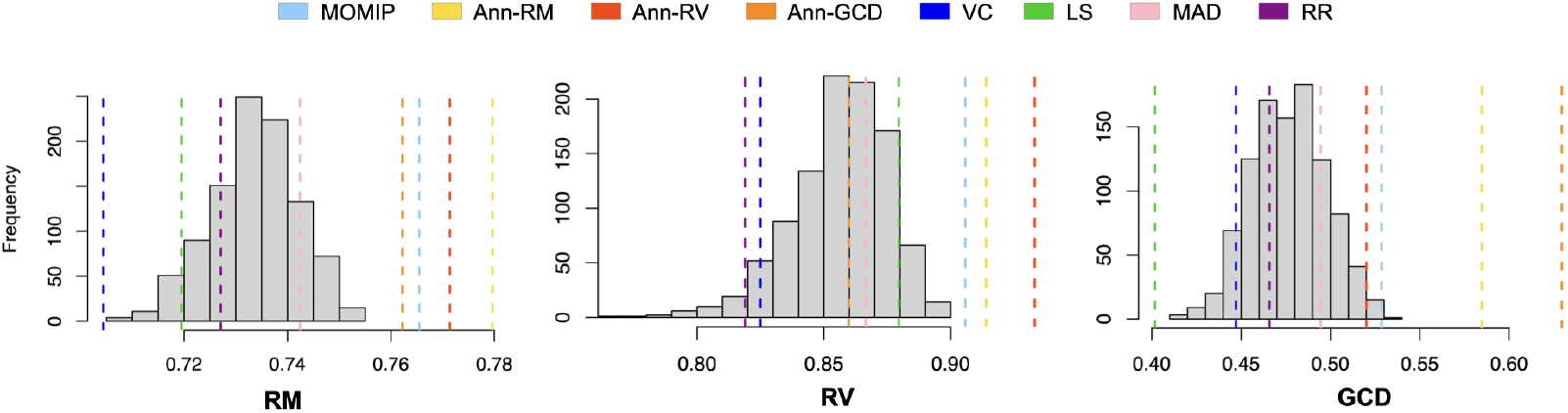
Representability coefficients for different unsupervised variable selection methods. The histograms show, from left to right, the RM, RV and GCD coefficients for 1000 random subsets of 18 variables of the full starting dataset. Vertical lines correspond to the coefficients computed by considering the subsets of variables identified by unsupervised variable selection methods, namely: MOMIP (light blue), Anneal based on RM, RV and GCD criteria (yellow, red, and orange, respectively), VC (blue), LS (green), MAD (pink), and RR (purple). Abbreviations: MOMIP, Max-Out Min-In problem; Ann-RM, Anneal based on RM; Ann-RV, Anneal based on RV; Ann-GCD, Anneal based on GCD; VC, Variance Criterion; LS, Laplacian Score; MAD, Mean Absolute Difference; RR, Relevance and Redundancy method.

**Figure 3:**
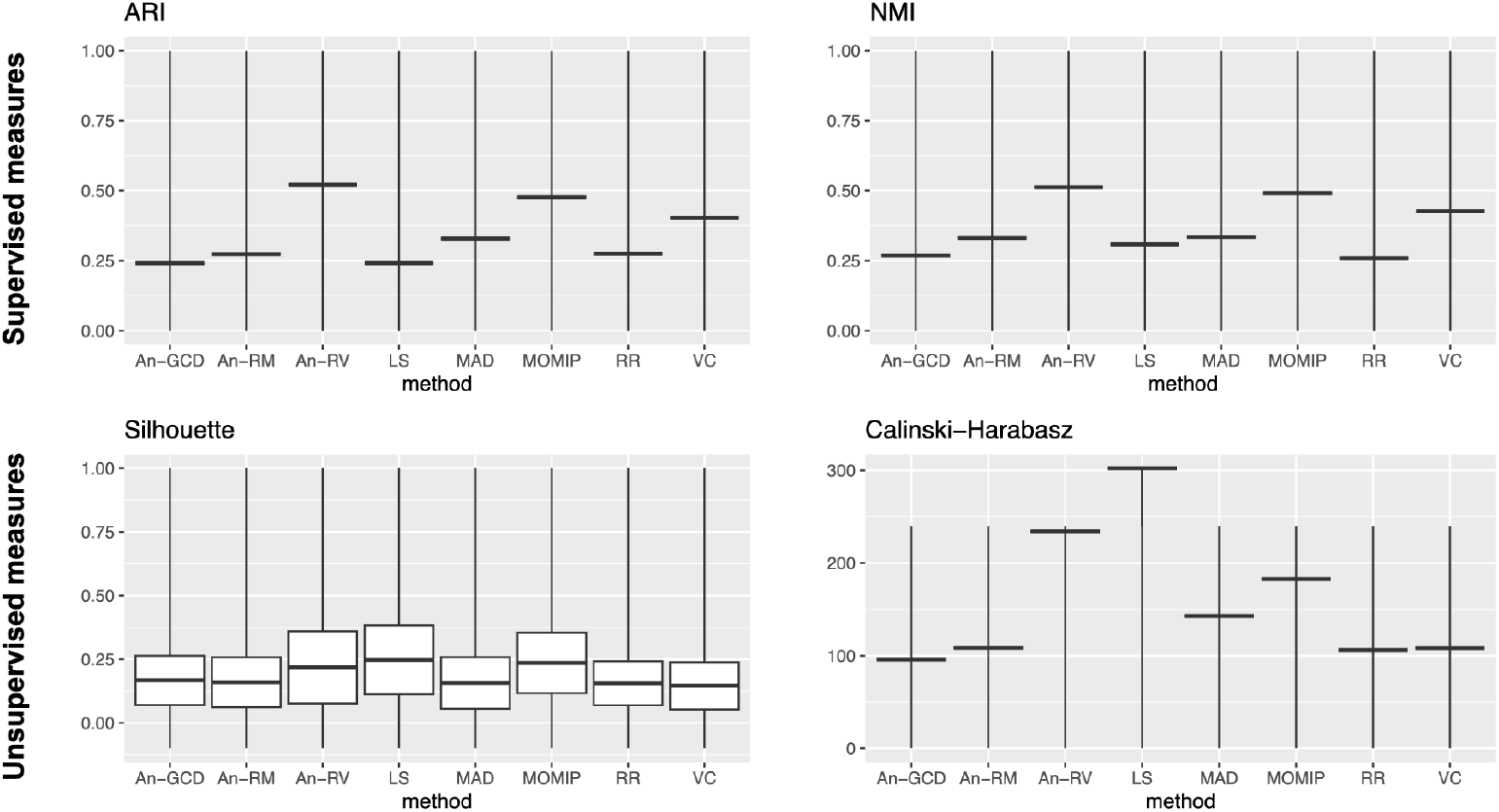
Performances of K-means clustering applied to dataset reduced to the subsets of 18 variables selected by different unsupervised methods. Supervised (ARI and NMI) and unsupervised (Silhouette and Calinski-Harabasz) measures are considered for evaluating clustering results. Scores are reported on y-axis, while methods are listed on the x-axis. Silhouette-related chart reports the mean value and the standard deviation over the silhouette values computed for each sample. Abbreviations: MOMIP, Max-Out Min-In problem; Ann-RM, Anneal based on RM; Ann-RV, Anneal based on RV; Ann-GCD, Anneal based on GCD; VC, Variance Criterion; LS, Laplacian Score; MAD, Mean Absolute Difference; RR, Relevance and Redundancy method.

As expected, the anneal method–which is specifically designed to find the optimal subset of 18 variables according to the RM, RV, or GCD criterion– led to the best results for the corresponding criteria. However, the outcomes appeared to be influenced by the choice of the criterion. For instance, the best performance was achieved with RM (yellow line), while using GCD (orange line) resulted in lower RV coefficients, similar to those obtained from random subsets. Overall, the MOMIP (light blue line), without parameter dependency, produced results comparable to the anneal method and was consistently ranked among the top-performing approaches. Conversely, the performance of the other unsupervised methods varied depending on the criterion used, with VC, LS, and RR yielding the worst results.

K-means clustering was used to evaluate how well the feature subsets, obtained by the variable-selection approaches, could recognize known glioma types in an unsupervised manner. The clustering performance was assessed using supervised measures (ARI, AMI, and NMI), which compare the identified clusters with the glioma types associated with each patient. Additionally, the quality of the clusters was evaluated using unsupervised silhouette and Calinski-Harabasz scores. The complete results are provided in Table S1 of the Supplementary Materials, while Figure offers a graphical summary of the performance for each criterion (note: the AMI coefficient is not shown because its results overlapped with those of NMI—refer to Table S1).

In agreement with the previous results on representability, the MOMIP and the anneal method (based on the RV criterion) demonstrated the best performance in recognizing the known classes, with moderate recovering of the glioma types (ARI, AMI, and NMI) around 0.5. However, the anneal method’s result was again influenced by the chosen criterion, and its effectiveness was significantly reduced when using RM or GCD. In fact, the anneal method based on GCD, along with LS and RR, was among the least effective in preserving the biological differences between glioma types. On the other hand, unsupervised measures revealed that the best overall clustering structure was obtained with the subset of variables identified by the LS approach, which achieved the highest average silhouette and Calinski-Harabasz scores. MOMIP also performed well, with a good Calinski-Harabasz score and a silhouette score comparable to LS, but with less variability across samples, suggesting a generally stable clustering structure.

These results confirm MOMIP as the most consistent method, yielding strong performance in both representability and the ability to preserve the biological differences and similarities) present in the original dataset.

### 3.2. TRIM-IT applied to GBM data

GBM is the most aggressive type of glioma. GBM patients can exhibit different tumor progressions, leading to diverse prognoses and suggesting the existence of subtypes that are not yet well defined. This study aims to identify a novel GBM stratification based on gene expression data using unsupervised K-means clustering. However, reducing the number of observations in the dataset (from 465 glioma patients to 116 GBM patients) creates an imbalance between variables and observations, which can lead to computational issues that may affect the results. To mitigate this, TRIM-IT can be used.

The first step of TRIM-IT consists in applying MOMIP to reduce the dimensionality of the initial dataset before applying clustering and conducting further analysis. Consistently with the workflow defined in the previous exploration phase, MOMIP was applied in two recursive steps, considering the dataset comprising 116 GBM samples and 145 variables, leading to a final selection of 19 variables.

Before applying K-means clustering (second step of TRIM-IT), we explored the GBM dataset to determine the optimal number of clusters *k*, a crucial step as the GBM subtypes are unknown. We considered different criteria for identifying the optimal *k*, namely, the silhouette score and the gap statistic, implemented by fviz nbclust R function [35]. Figure shows the results obtained for *k* ranging from 1 to 10 using the two criteria. The vertical lines indicate the optimal number of clusters, which differs between the two methods. The highest silhouette score is achieved at *k* = 3, while gap statistics suggests *k* = 4 as the preferable choice. While setting *k* = 4 may capture marginal differences, the more conservative choice of *k* = 3 ensures compact clusters and facilitates meaningful biological insights. Therefore, we have selected *k* = 3 as the most reliable option.

K-means clustering was then applied to both the GBM dataset (116 samples and 145 variables) and the MOMIP-reduced one (116 samples and 19 variables). When comparing the two clustering results (Table 1), we observe that many samples are allocated to the same clusters, with some mislabeling, particularly between clusters 1 and 3.

**Table 1:** Cross-comparison among clusters defined by K-means applied to the initial GBM dataset (rows) and the MOMIP-reduced one (columns). Each cell reports a number corresponding to the samples assigned to the corresponding cluster; diagonal cells highlight the agreement between the two results, representing the number of samples assigned to the same cluster in the two cases. Abbreviations: DS, dataset; MOMIP, Max-Out Min-In Problem; C1, Cluster 1; C2, Cluster 2; C3, Cluster 3.

|  |  | MOMIP-reduced DS |  |  |
| --- | --- | --- | --- | --- |
|  |  | C1 | C2 | C3 |
| GBM DS | C1 | 41 | 2 | 21 |
|  | C2 | 0 | 21 | 9 |
|  | C3 | 4 | 1 | 17 |

Table reports the clustering performance in the two cases. The higher silhouette and Calinski-Harabasz scores obtained after MOMIP application confirm a better clustering structure with the reduced dataset. To further support this result, the GBM patients were visualized by the Uniform Manifold Approximation and Projection (UMAP) technique for non-linear dimension reduction [36]. Figure shows the projection of the samples in a two-dimensional space based on the gene expression data from (a) the initial GBM dataset and (b) the MOMIP-reduced dataset, with colors indicating the identified clusters. Figure 5(a) exhibits some overlap, whereas Figure 5(b) shows a clearer separation between the groups. Particularly, clusters 1 and 3 are well-separated along the y-axis, while cluster 2 is distinct along the x-axis, forming three clearly defined groups in the two-dimensional space. These results suggest that the MOMIP method not only effectively reduces dimensionality but also enhances the quality of the identified clusters.

**Figure 4:**
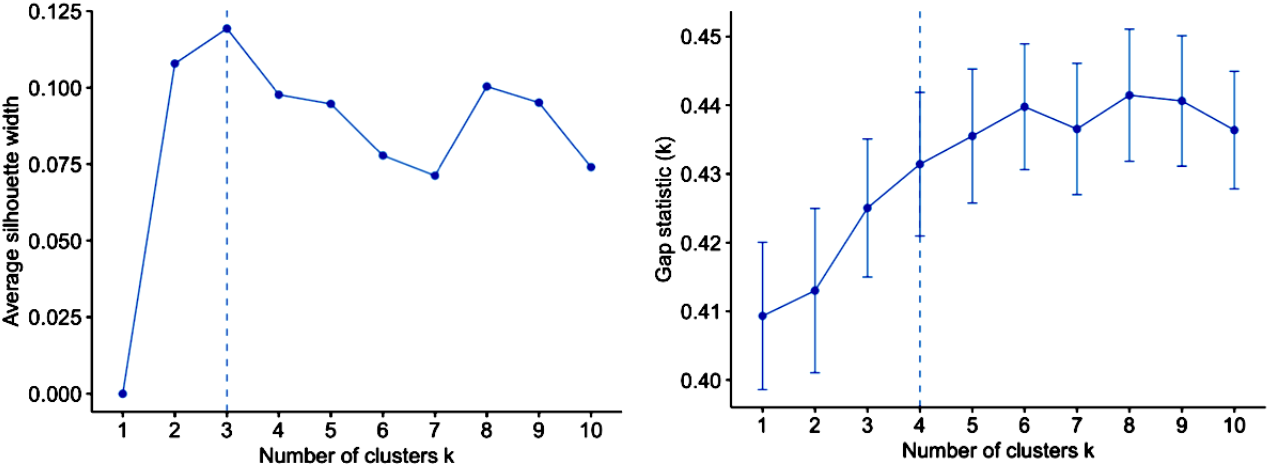
Optimal number of clusters for GBM data. Charts show the results depending on the number of clusters (*k*) of different criteria, namely, from left to right, silhouette,and gap statistic. Dotted lines correspond to the optimal value for each criterion. Results are computed by the fviz nbclust R function.

**Figure 5:**
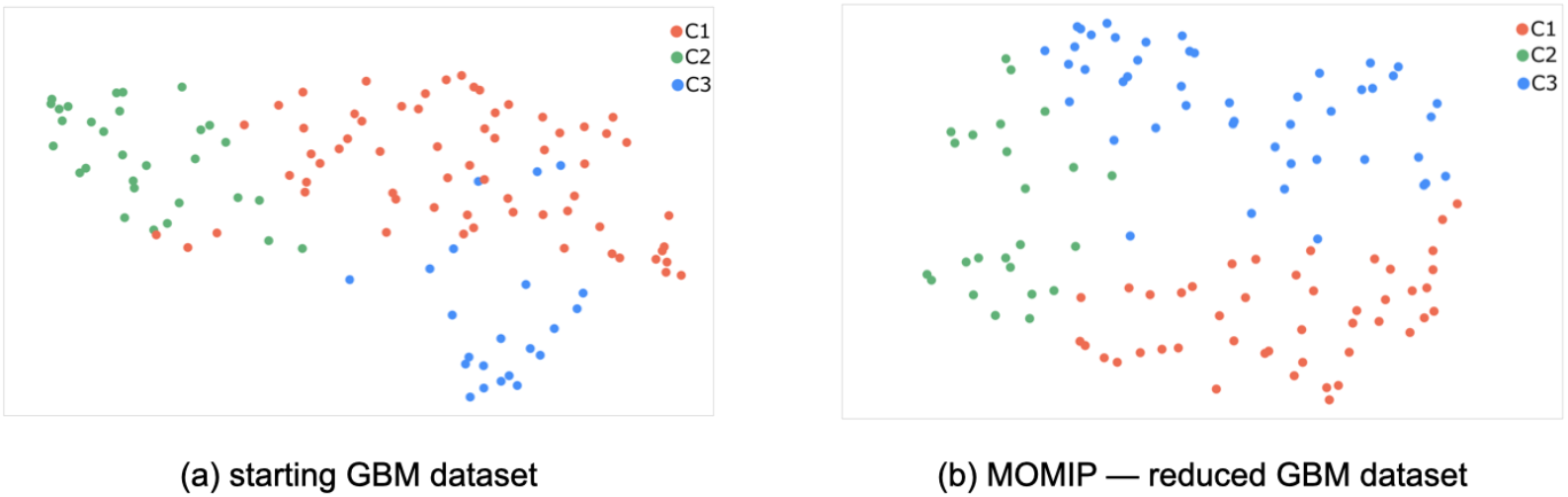
UMAP of GBM data by taking into account (a) the starting GBM dataset, and (b) the reduced dataset with the subset of variables selected by MOMIP. Clusters identified by K-means clustering are highlighted with different colors: red, Cluster 1 (C1); green, Cluster 2 (C2); blue Cluster 3 (C3).

**Figure 6:**
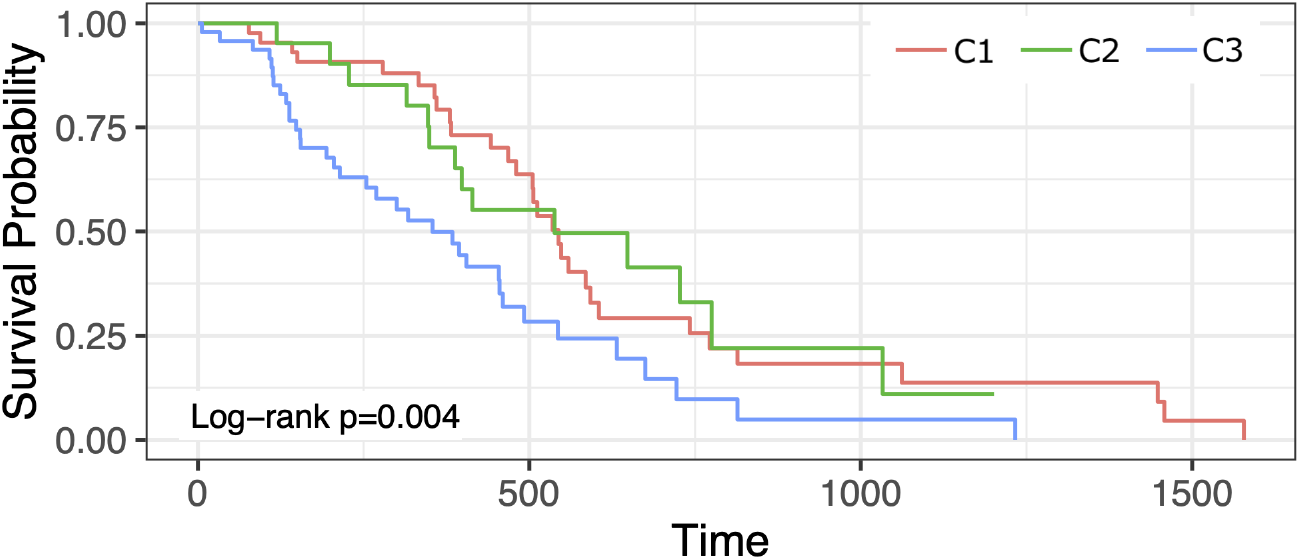
Kaplan-Meier curves on the overall survival of the patients of each of the three GBM clusters identified by K-means clustering (log-rank p-value=0.004). Red, Cluster 1 (C1); green, Cluster 2 (C2); blue Cluster 3 (C3).

**Figure 7:**
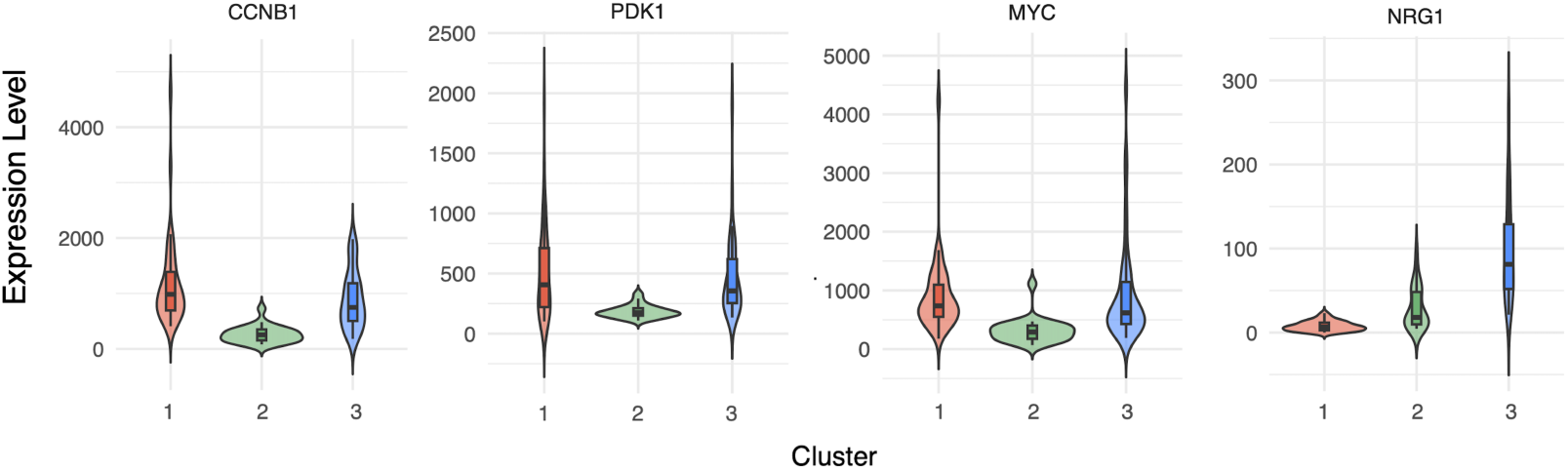
Top four genes with higher differential expression among the three groups. Gene expression level over each cluster is reported on y-axis. Clusters identified by K-means clustering are highlighted with different colors: red, Cluster 1; green, Cluster 2; blue Cluster 3. Complete results of the differential gene expression analysis are available as Supplementary Materials section S3

After verifying that the identified clusters are meaningful from a technical perspective, we aim to investigate whether there are properties that define each cluster. Specifically, our goal is to determine whether there are distinct features that characterize the samples within each cluster, potentially identifying a GBM subtype.

First, we examined whether there was any correlation between these clusters and the available patients’ clinical information (which can be retrieved from TCGA alongside the RNA-Seq data). The clinical features considered for potential correlation with the patient groups included gender, radiation therapy status, race, and histology. Since both radiation therapy and race data were too imbalanced to yield statistically significant results, only gender and histology were analyzed.

We assessed the correlation between these clinical features and the three clusters using the Chi-Square test, which determines if there is a significant relationship between two categorical variables, and the Cramer V index, which quantifies the strength of the correlation. The analysis revealed no correlation between gender and the identified clusters. However, there was a statistically significant association between clusters and histology, with a Chi-Square value of 29.61 (p-value = 3.72e-07) and a Cramer’s V index of 0.51, indicating a strong relationship.

Histology provides information about the microscopic structure of tumor tissue. Gliomas can vary in cellular composition, and can present distinct characteristics depending on the severity of the cancer condition (tumor grade). Typically, GBM is characterized by a peculiar high-grade histology. Nevertheless, there are some GBM patients that display histology typical of Lower-Grade Glioma (LGG), while still having the molecular features of GBM. Interestingly, an examination of the histology of the observations in each cluster (see Table 3) shows that cluster 2 appears to be characterized by LGG-like histology, while clusters 1 and 3 predominantly consist of samples with typical GBM features.

**Table 2:** Comparison of clustering performance of the full GBM dataset comprising 145 variables), and the MOMIP-reduced dataset (with 19 variables). Performance is evaluated by unsupervised criteria, namely average silhouette value and Calinski-Harabasz score.

| Scores | Full GBM DS | MOMIP-reduced |
| --- | --- | --- |
| Average Silhouette | 0.12 | 0.16 |
| Calinski-Harabasz | 15.13 | 24.68 |

**Table 3:** Cross-comparison among GBM clusters (rows) and samples’ histology (columns). Each cell reports a number corresponding to the samples assigned to the clusters on the rows, having the histology defined in the columns. Abbreviations: LGG, histology typical of Lower-Grade Glioma; GBM, histology typical of glioblastoma; C1, Cluster 1; C2, Cluster 2; C3, Cluster 3.

|  | Histology |  |
| --- | --- | --- |
|  | GBM | LGG |
| C1 | 33 | 12 |
| C2 | 2 | 22 |
| C3 | 31 | 16 |

The clinical GBM data also include information on the patient survival status, enabling us to perform a survival analysis to identify potential differences in life expectancy between patients in each cluster. The Kaplan-Meier curves of the three clusters in Figure reveal a statistically significant difference in survival (p-value = 0.004). Specifically, cluster 3 is associated with poor overall survival, whereas clusters 1 and 2 show better prognosis. To further investigate the significance of survival differences between cluster pairs, we first computed the pairwise log-rank p-value with Benjamini-Hochberg adjustment [37]. This analysis revealed that clusters 1 and 2 are not significantly different (p-value = 0.9131), while clusters 1 and 3 have statistically significant difference (p-value=0.0095). When comparing clusters 3 and 2, the log-rank p-value was close to the 0.05 significance threshold (p-value = 0.052), suggesting the need for further analysis. To this end, Cox regression was applied, treating the assigned clusters as categorical variables.The analysis confirmed the high similarity between clusters 1 and 2, with a hazard ratio (HR) of 1, while highlighting a significant difference (p-value = 0.017) between clusters 2 and 3. The HR of 2,15 suggests that cluster 3 has approximately twice the risk of death compared to cluster 2. These results are provided in Supplementary Materials S2, Table S3.

The next step of our study focuses on examining the molecular profile to identify genes that are differentially expressed across the three clusters and may serve as potential biomarkers. We conducted Differential Gene Expression (DGE) analysis by edgeR R package[38] (details provided in the Supplementary Materials section S3).

The results revealed several genes differentially expressed across the three clusters. The complete results of the DGE analysis are reported in Supplementary Tables S4, S5, and S6, while the Figure summarizes the mean expression level of the top four differentially expressed genes. Cluster 2 was characterized by reduced expression of several genes compared to clusters 1 and 3, including *CCNB1, PDK1*, and *MYC*.

Interestingly, these genes are typically overexpressed in GBM and are associated with tumor cell proliferation and cancer progression [39, 40, 41], suggesting that patients in cluster 2 exhibit an atypical molecular profile compared to most GBM cases. Notably, silencing these genes has demonstrated therapeutic benefits, inhibiting tumor growth and improving prognosis in bladder cancer [42] and glioma [41, 43, 40].

Although clusters 1 and 3 appear to be molecularly similar, they exhibit a significant difference in the expression of the *NRG1* gene. This gene is highly overexpressed in cluster 3 and may therefore be associated with worse survival. This finding is supported by previous studies investigating *NRG1* as a GBM biomarker, showing its correlation with tumor aggressiveness and poor prognosis [44, 45, 46].

## 4. Conclusions

This study focuses on the application of unsupervised variable selection methods to glioma gene expression data. Variable selection is particularly beneficial for high-dimensional datasets where the number of features exceeds the number of observations, and no prior knowledge exists to guide a supervised selection strategy. While variable selection can be effectively implemented into methods aimed at specific tasks (such as survival analysis or clustering), performing *a priori* unsupervised selection proves invaluable for reducing dimensionality without being tied to any particular goal. This approach minimizes the risk of model overfitting, ensuring that the data’s underlying structure and information are preserved. The resulting reduced dataset can then be used across a range of analyses, maintaining flexibility and applicability without task-specific constraints.

We evaluated the performances of several methods on glioma RNA-Seq data, and identified MOMIP as the most consistent one, making it suitable for our purposes. Our ultimate goal was to employ unsupervised variable selection to conduct further studies on GBM, the most aggressive glioma with the poorest prognosis. To do this, we defined a new tool called TRIM-IT, structured in four consequent steps. First, MOMIP was applied to the GBM dataset to select 19 variables. The reduced dataset was then used for patient stratification via K-means clustering, resulting in well-defined clusters. These groups were subsequently analyzed to uncover clinical features that characterized the patients in each group, including survival data. Finally, gene expression data were analyzed to identify genes that were differentially expressed across the clusters through DGE analysis.

The examination of the clinical profile of the patients revealed distinct traits: cluster 2 was associated with LGG-like histology, and cluster 3 was characterized by poor survival outcomes. Moreover, DGE analysis revealed that cluster 2 exhibited lower expression levels of many genes, while cluster 1 and 3 were discriminated by low and high expression of the gene *NRG1*, respectively, which might be associated with poor prognosis, and therefore it is a good candidate as a prognostic biomarker.

There are some limitations to our study. First, we considered a prefiltered RNA-Seq dataset that retains genes associated with major cancerrelated processes but excludes other variables that could be important for more comprehensive analyses. This dataset reduction was implemented as MOMIP was solved using quadratic unconstrained binary optimization, which, while providing an exact solution, is constrained by the data’s dimensionality. To handle larger datasets, MOMIP would need to be solved by using an optimization algorithm for approximate solutions, such as anneal. As a next step, we plan to modify the MOMIP implementation by incorporating the anneal algorithm, enabling it to process more extensive datasets effectively. Second, the two-step variable selection that we performed aimed to reduce the variables to a relatively small amount that might be easily analyzed; depending on the data considered, this approach may need to be adjusted. Third, data normalization can impact the performances of unsupervised variable selection methods. If it is necessary (as in our case) data transformations preserving the overall data variability should be preferred.

In conclusion, this work provides an overview of unsupervised variable selection methods and introduces a framework for evaluating their performance based on the data characteristics to preserve. Furthermore, we present a new tool that performs unsupervised pre-selection to identify key variables for patient stratification, making them potential diagnostic and/or prognostic biomarkers. Our analysis pointed out the potential of the MOMIP method, particularly suitable in cases where avoiding parameter dependency is a priority. Moreover, our investigation of the GBM dataset via TRIM-IT revealed three clusters with strong potential to define GBM subtypes. Further studies are needed to biologically validate our findings and confirm the clinical relevance of the identified subgroups.

## CRediT author statement

**Roberta Coletti:** Conceptualization, Methodology, Software, Validation, Formal analysis, Data Curation, Visualization, Writing - Original Draft. **J. Oreste Cerdeira:** Conceptualization, Methodology, Validation, Writing - Original Draft, Supervision. **Marcos Raydan:** Conceptualization, Methodology, Validation, Writing - Review & Editing, Supervision. **Marta B. Lopes:** Conceptualization, Methodology, Validation, Writing - Review & Editing, Supervision, Funding acquisition.

## Acknowledgements

This work was supported by national funds through Fundação para a Ciência e a Tecnologia (FCT) with references CEECINST/00042/2021, UIDB/00297/2020 and UIDP/00297/2020 (NOVA Math, Center for Mathematics and Applications), UIDB/00667/2020 and UIDP/00667/2020 (UNIDEMI), and the research project PTDC/CCI-BIO/4180/2020 “MONET – Multi-omic networks in gliomas” (doi: 10.54499/PTDC/CCI-BIO/4180/2020). The results presented here are based upon data generated by the TCGA Research Network: https://www.cancer.gov/tcga.

