## Supplementary Materials for "Performance Assessment of an Unsupervised Variable Selection Approach for Biomarker Discovery and Glioblastoma Subtyping"

This document contains supplementary contents that complements and supports the analysis presented in the main text.

#### S1 Clustering

Clustering was used to evaluate whether the variable selection process negatively impacted patient grouping, using the clustering from the full glioma dataset as a reference. Table S1 summarizes the results.

Table S1: Comparison of clustering results based on glioma datasets containing the same samples but different sets of variables. The gray column represents the full glioma dataset, which includes all samples (206 astrocytomas, 143 oligodendrogliomas, and 116 GBM) and 145 bulk gene expression variables, which serves as a reference. All other columns refer to reduced datasets with only 18 selected variables, which vary depending on the method used. The methods include the Max-Out Min-In Problem (MOMIP); anneal optimization based on RM (Ann-RM), RV (Ann-RV), or GCD (Ann-GCD) criteria; Variance Criterion (VC); Laplacian Score (LS); Mean Absolute Difference (MAD); Relevance and Redundancy (RR). Clustering performance is evaluated using Adjusted Rand Index (ARI), Adjusted Mutual Information (AMI), Normalized Mutual Information (NMI), Average Silhouette (Sil), and Chalinski-Harabasz score (CH).

| Scores | Full DS | MOMIP | Ann-RM | Ann-RV | Ann-GCD | VC | LS | MAD | RR |
|---|---|---|---|---|---|---|---|---|---|
| ARI | 0.60 | 0.48 | 0.27 | 0.52 | 0.24 | 0.40 | 0.24 | 0.33 | 0.28 |
| AMI | 0.57 | 0.49 | 0.33 | 0.51 | 0.27 | 0.42 | 0.30 | 0.33 | 0.26 |
| NMI | 0.58 | 0.49 | 0.33 | 0.51 | 0.27 | 0.43 | 0.31 | 0.33 | 0.26 |
| Sil | 0.12 | 0.24 | 0.16 | 0.22 | 0.17 | 0.15 | 0.25 | 0.16 | 0.16 |
| CH | 87.52 | 182.62 | 108.66 | 234.14 | 95.95 | 108.38 | 302.31 | 142.84 | 106.11 |

#### S2 Survival Analyisis

Survival analysis was performed on the three clusters identified from the MOMIP-reduced GBM dataset. To assess whether the differences between the Kaplan-Meier curves were statistically significant, the pairwise log-rank p-values were computed using the `pairwise_survdiff` function from the `survminer` R package. Given that multiple comparisons were made, the Benjamini-Hochberg method–already implemented in the function–was used to adjust the p-values. The results are provided in the table S2.

Table S2: Pairwise log-rank p-values for the survival distribution of the GBM's clusters. C1, cluster 1; C2, cluster 2; C3, cluster 3.

| | C1 | C2 |
|---|---|---|
| C2 | 0.9131 | – |
| C3 | 0.0095 | 0.0517 |

To further analyze the difference in survival between clusters 2 and 3, Cox regression was performed with cluster 2 as the reference. Then, `coxph` function from `survival` R package was used to fit a Cox proportional hazards regression model. The resulting Hazard Ratio (HR), the relative confidence interval (CI), and the corresponding p-value are shown in Table S3.

Table S3: Survival cluster comparison by Cox regression. Rows refer to the two cases of study "Cluster 1 vs Cluster 2" (C1 vs C2) and "Cluster 2 vs Cluster 3" (C2 vs C3). Columns report the Hazard Ratio (HR), with the corresponding lower and upper 95% Confidence Interval (CI) and p-value.

| | HR | Lower 95% CI | Lower 95% CI | p-value |
|---|---|---|---|---|
| C1 vs C2 | 0.9997 | 0.5227 | 1.912 | 0.9992 |
| C2 vs C3 | 2.1451 | 1.1452 | 4.018 | 0.0172 |

### S3 Differential Gene Expresion Analysis

For Differential Gene Expression (DGE) analysis, we performed quasi-likelihood (QL) F-test by the `EdgeR` R package. Considering the raw data and the cluster assigned each patient, `glmQLFit` function fits a quasi-likelihood negative binomial generalized linear model (GLM) to the gene expression data. This model is then used to perform the F-test by the glmQLFTest R function, which allows for comparison between cluster pairs. The top 10 genes for each pair of clusters are resumed in Tables S4, S5, and S6.

Genes are ranked based on the F-statistic (F), which evaluates whether adding the cluster variable significantly improves the model's fit. Higher F statistic indicates stronger evidence that gene expression differs between clusters. The statistical significance of this difference is assessed using the p-value, which is adjusted for multiple testing using the false discovery rate (FDR); an FDR below 0.05 is commonly used as a significance threshold.

The function also returns the log Fold Change (logFC), which quantifies the difference in gene expression between the two clusters on a logarithmic scale. A positive logFC value means the gene is downregulated in the first group compared to the second, while a negative logFC value means it is upregulated. Additionally, the tables include the log Counts Per Million (logCPM), which represents the average expression level of the gene across all samples.

Table S4: Top 10 genes resulted from the Differentially Gene Expression analysis of cluster 1 vs cluster 2. Gray cells highlight statistically significant genes.

| Gene | logFC | logCPM | F | PValue | FDR |
|---|---|---|---|---|---|
| *CCNB1* | -1.9260675 | 14.21512 | 80.102037 | 5.731907e-15 | 1.089062e-13 |
| *NRG1* | 2.0665280 | 10.14007 | 55.906415 | 1.422814e-11 | 1.351673e-10 |
| *PDK1* | -1.3735788 | 13.22406 | 33.426954 | 6.085314e-08 | 3.854032e-07 |
| *PRDX1* | 0.5211548 | 17.45013 | 30.249629 | 2.210354e-07 | 1.049918e-06 |
| *MYC* | -1.3889054 | 14.11526 | 29.573836 | 2.919794e-07 | 1.109522e-06 |
| *RICTOR* | 0.6478297 | 13.61395 | 20.927843 | 1.178748e-05 | 3.732704e-05 |
| *MAPK3* | 0.3325375 | 15.26139 | 12.773545 | 5.089917e-04 | 1.381549e-03 |
| *XRCC5* | -0.2249615 | 16.87397 | 9.521406 | 2.526650e-03 | 6.000794e-03 |
| *MTOR* | 0.2312891 | 15.20338 | 3.357087 | 6.941883e-02 | 1.465509e-01 |
| *ERRFI1* | -0.4565118 | 14.48526 | 3.178998 | 7.714208e-02 | 1.465700e-01 |

Table S5: Top 10 genes resulted from the Differentially Gene Expression analysis of cluster 1 vs cluster 3. Gray cells highlight statistically significant genes.

| Gene | logFC | logCPM | F | PValue | FDR |
|---|---|---|---|---|---|
| *NRG1* | 3.6938170 | 10.14028 | 181.264791 | 1.161497e-25 | 2.206844e-24 |
| *RPS6KA1* | 0.3560954 | 14.20658 | 6.433393 | 1.249588e-02 | 1.187108e-01 |
| *PRDX1* | 0.1765165 | 17.45013 | 4.927251 | 2.833366e-02 | 1.517771e-01 |
| *CCNB1* | -0.3446113 | 14.21529 | 4.520364 | 3.556276e-02 | 1.517771e-01 |
| *NOTCH1* | -0.3313189 | 16.40741 | 4.314588 | 3.994133e-02 | 1.517771e-01 |
| *RICTOR* | 0.2270492 | 13.61394 | 3.623907 | 5.937328e-02 | 1.880154e-01 |
| *ERRFI1* | 0.3259023 | 14.48548 | 2.453725 | 1.199052e-01 | 3.254569e-01 |
| *AKT1S1* | -0.1473139 | 14.73894 | 2.048819 | 1.549488e-01 | 3.508274e-01 |
| *MAPK3* | 0.1080440 | 15.26142 | 1.940806 | 1.661814e-01 | 3.508274e-01 |
| *PDK1* | -0.1927302 | 13.22458 | 1.093680 | 2.977797e-01 | 5.657814e-01 |

Considering only the top genes reported in Tables S4, S5, and S6 with FDR$<0.05$, Figure S1 shows their mean expression level across clusters.

Table S6: Top 10 genes resulted from the Differentially Gene Expression analysis of cluster 2 vs cluster 3. Gray cells highlight statistically significant genes.

| Gene | logFC | logCPM | F | PValue | FDR |
|---|---|---|---|---|---|
| *CCNB1* | 1.5813104 | 14.21529 | 56.561211 | 1.137174e-11 | 2.160631e-10 |
| *MYC* | 1.3841392 | 14.11534 | 29.671119 | 2.806805e-07 | 2.666465e-06 |
| *NRG1* | 1.6321807 | 10.14028 | 26.353601 | 1.123307e-06 | 7.114278e-06 |
| *PDK1* | 1.1811059 | 13.22458 | 25.468975 | 1.636124e-06 | 7.771589e-06 |
| *PRDX1* | -0.3446324 | 17.45013 | 13.312314 | 3.929098e-04 | 1.493057e-03 |
| *XRCC5* | 0.2343777 | 16.87397 | 10.477261 | 1.566218e-03 | 4.959692e-03 |
| *ERRFI1* | 0.7829362 | 14.48548 | 9.180094 | 3.002565e-03 | 8.149820e-03 |
| *RICTOR* | -0.4210131 | 13.61394 | 8.879723 | 3.497417e-03 | 8.306366e-03 |
| *MAPK3* | -0.2244401 | 15.26142 | 5.885563 | 1.676844e-02 | 3.540005e-02 |
| *MTOR* | -0.2842826 | 15.20332 | 5.174979 | 2.470727e-02 | 4.694382e-02 |

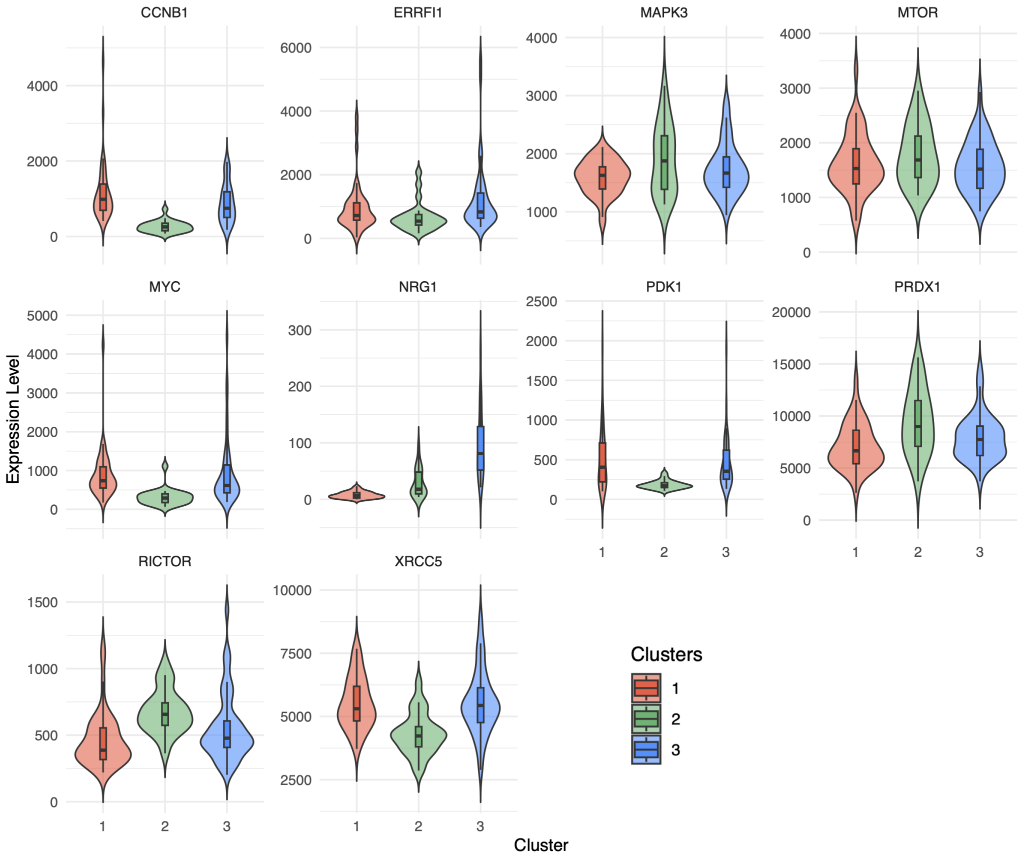


Figure S1: Comparison of gene expression levels across clusters via violin plot. Each chart refers to a specific gene, where clusters are highlighted with different colors: red, Cluster 1; green, cluster 2; blu, cluster 3.
